# Afterimages drive a shared visual motion-reversal illusion in *Drosophila*

**DOI:** 10.64898/2026.01.19.700413

**Authors:** Heng Wu, Tong Gou, Damon A. Clark

## Abstract

Illusions expose core computations in perception. In one visual apparent-motion illusion, perceptual direction is reversed when phase-shifted gratings are interleaved with uniform frames. Here, we demonstrate that *Drosophila* exhibits the same direction reversal reported in mammals. Combining behavior, targeted silencing, two-photon imaging, and modeling, we localize the origin of this illusion to elementary motion pathways. Silencing direction-selective T4/T5 neurons abolishes the reversal, and recordings reveal that downstream wide-field neurons invert their directional preference as interleave duration increases. Replacing periodic gratings with random binary patterns preserves the reversal, implicating afterimages rather than spatial periodicity. Imaging neurons upstream of T4/T5 shows signatures of an afterimage, whose emergence depends on interleave luminance. Critically, dark interleaves suppress afterimages and eliminate both the neural and behavioral reversal, whereas light interleaves preserve or enhance it. Thus, afterimages are central to this shared illusion and explain a deficiency of canonical motion-energy accounts. These results link a classic apparent-motion phenomenon to identified circuit elements and reveal a simple stimulus manipulation that switches an illusion on and off.

## Introduction

Sometimes, visual stimuli elicit percepts that deviate from expectations, a phenomenon commonly referred to as an illusion (Gregory, 1997; Todorović, 2020). Illusions can often provide insight into the principles and mechanisms underlying visual processing (Eagleman, 2001). For instance, Mach band illusions illustrate how the center-surround organization in the visual system impacts perception (Lotto et al., 1999; Syrkin et al., 1994), while reverse-phi illusions emphasize the reliance of motion detection algorithms on pairwise correlations (Anstis, 1970; Anstis & Rogers, 1986; Bialek & van Steveninck, 2005; Eichner et al., 2011; Hassenstein & Reichardt, 1956; Leonhardt et al., 2017; Salazar-Gatzimas et al., 2018; Salazar-Gatzimas et al., 2016; Tuthill et al., 2011). Among these, illusions shared across divergent species or multiple sensory modalities are particularly compelling (Agrochao et al., 2020; Allik et al., 1989; Bååth et al., 2014; Lynna C. Feng et al., 2017; L. C. Feng et al., 2017; Gori et al., 2014; Kadakia et al., 2022; Orger et al., 2000; Vaziri et al., 2024; Wu et al., 2020), as they highlight visual processing potentially shared across phyla. Such shared percepts suggest deep similarities in perceptual strategies and sensory algorithms (Borst & Helmstaedter, 2015; Clark & Demb, 2016).

In this study, we leverage a motion reversal illusion to explore mechanisms of motion detection—a critical ability enabling animals to detect object motion and optical flow, essential for perceiving and interacting with their environment (Borst et al., 2010; Nishida, 2011; Rokszin et al., 2010; Wei, 2018; Yang & Clandinin, 2018).

Human observers experience an illusory motion reversal when uniform gray frames are interleaved between displacements of a sinusoidal grating (Pantle & Turano, 1992; Sheliga et al., 2006; Strout et al., 1994; Takeuchi & De Valois, 1997; Takeuchi et al., 2001). In these experiments, human participants were presented with a stimulus consisting of sequential stationary sinusoidal contrast gratings, each shifted by some phase. Because the stimulus is periodic, these stimuli are fundamentally ambiguous about potential underlying motion, since many different phase shifts could correspond to the observed displacement. In humans, this ambiguity is resolved to generate a percept in the direction of the smallest shift of the sinusoid. However, if an interleaving frame of uniform intermediate luminance is introduced between the shifted gratings (**Fig. 1A**) (which we call a sinusoidal interleave-shift stimulus), then strikingly, humans often perceive a reversal of the motion direction (Pantle & Turano, 1992; Strout et al., 1994; Takeuchi & De Valois, 1997), so that motion is in the direction of the second shortest possible displacement. This reversal depends on the duration of the interleaved frames, so that interleaves of ∼50 ms tend to result in the strongest reversals.

**Figure 1.**
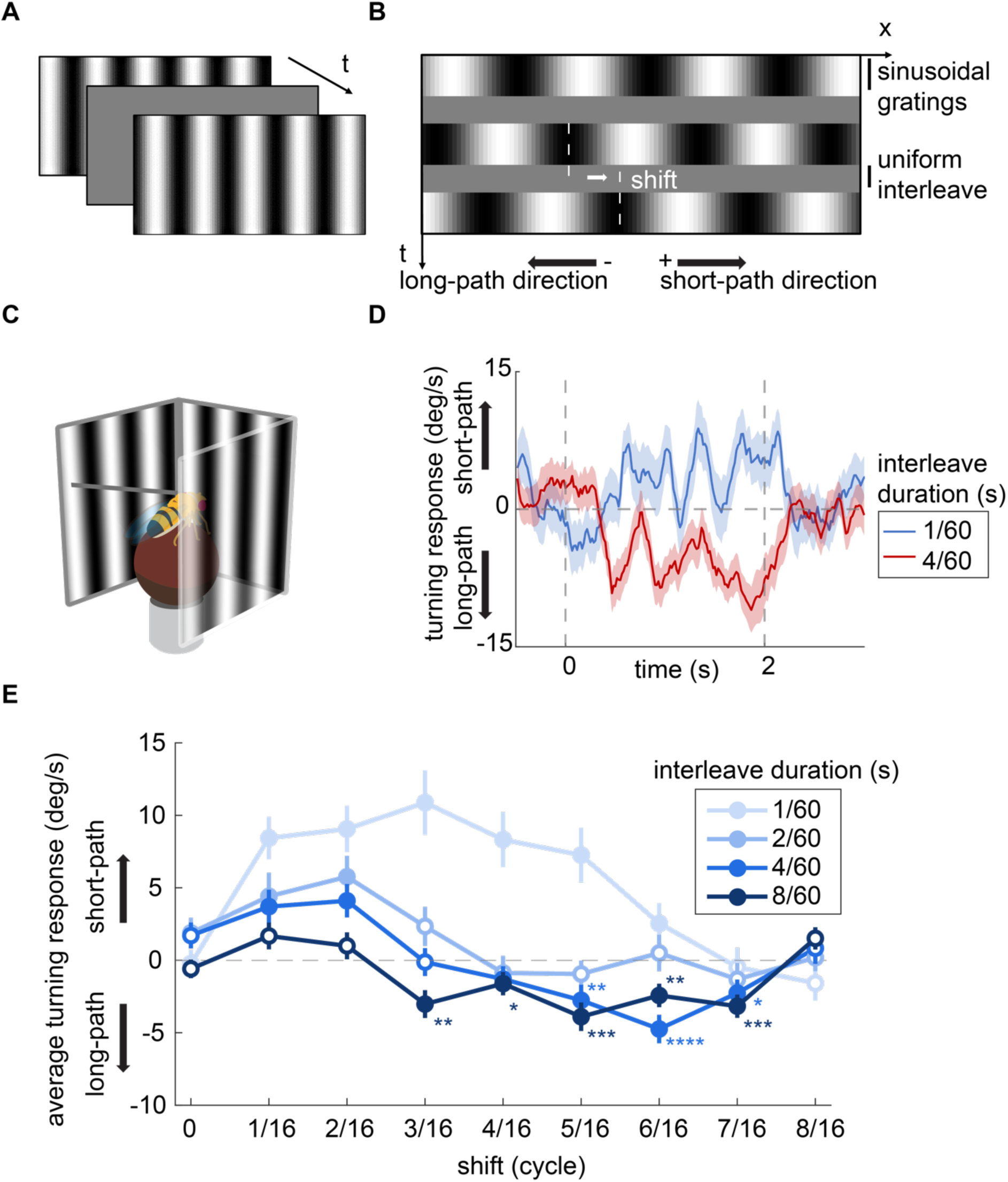
Flies perceive illusory motion-reversal for shifting sinusoid gratings with long interleaves. (A) Diagram of the illusions in human experiments, shifted sinusoid gratings interleaved by a uniform interleave. (B) Space-time contrast diagram of a typical stimulus in fly experiments: stationary full-contrast sinusoid gratings with 30° wavelength were displayed for 12/60 s, interleaved with uniform frames of full-field contrast. (C) In fly behavioral experiments, tethered flies were placed above air-supported balls, their turning response was measured when the stimuli were projected to the screens. (D) Typical time traces of wild-type fly turning behavior towards stimuli with different interleave durations, the shifts of both curves are 3/8 cycle. (E) Averaged turning response of flies towards stimuli with different shifts and interleave durations (N = 43, 33, 56, 56 flies for interleave duration = 1/60, 2/60, 4/60, 8/60 s respectively.). Error bars indicate mean ± SEM and the dot is filled if p < 0.05 by Wilcoxon signed-rank tests against 0. * p < 0.05, ** p < 0.01, *** p < 0.001and **** p < 0.0001 by Wilcoxon signed-rank tests against 0, shown only for negative significant values.

Subsequent research demonstrated that mice and monkeys also showed reversed optokinetic responses in response to sinusoidal interleave-shift stimuli (Miura et al., 2018; Takemura et al., 2021). This suggests that the underlying processing resulting in this reversal may be conserved across mammals, providing an opportunity to investigate shared principles of motion detection and perceptual disambiguation.

Previous studies also showed that a motion energy model (Adelson & Bergen, 1985) could account for the perceptual reversal in the interleave-shift stimulus under some conditions (Pantle & Turano, 1992; Strout et al., 1994; Takeuchi & De Valois, 1997). These analyses proposed that for longer interleave durations, the weighted energy in the reversed motion direction may exceed that of the motion in the direction of the smallest shift, leading to a reversed motion percept. However, this mathematical explanation has not been connected to response properties at the neuronal level.

In this study, we investigate the motion percept induced by the interleave-shift stimulus in the fruit fly *Drosophila*, a species evolutionarily distant from mammals (Kumar et al., 2022).

Fruit flies offer significant advantages as a model organism due to their well-mapped neural circuits (Dorkenwald et al., 2024; Scheffer et al., 2020; Schlegel et al., 2024) and the availability of powerful genetic tools (Luo et al., 2018). These characteristics allow us to dissect the neural processing driving this illusion, which is not easily accessible in mammalian models.

Here, we measured behavioral responses of *Drosophila* fruit flies to interleave-shift stimuli. We found that like mammals, flies resolve the ambiguity in the stimulus in favor of the shortest displacement when interleaves are absent; also like mammals, flies perceive reversed motion as the interleave increased in duration. By silencing direction-selective neurons, we demonstrated that this motion reversal is mediated by the direction-selective local motion detecting neurons, T4 and T5. Furthermore, by recording neural responses in T4 and T5 and in a downstream, widefield neuron, we found that their activity could explain the perceptual reversal observed in the fly’s behavior. In these circuits, the motion reversal can be explained by integrating signals from T4 and T5 neurons with different preferred directions, revealing the neuronal basis of the illusion. Theoretical results, psychophysical tests, and imaging of neurons upstream of motion detectors all suggested that afterimages played a central role in the perceptual reversal. We discovered we could eliminate afterimages under some interleave conditions, and in those cases, abolish the illusory reversal, demonstrating that afterimages are critical for generating this illusory motion reversal. Overall, these results reveal the neuromechanistic origins of this illusion in fruit flies and the importance of afterimages in generating this apparent-motion reversal.

## Results

### Flies reverse turn direction in response to the interleave-shift stimuli

We first asked whether flies, like humans and mice, exhibit direction reversals in their perceptual responses to the sinusoidal interleave-shift stimulus. We developed a stimulus paradigm for flies in which sinusoidal gratings were continuously shifted and interleaved with uniform, mean luminance (**Fig. 1A-B**). Flies were tethered and placed above air-supported balls while stimuli were projected onto panoramic screens around them (**Fig. 1C**) (Creamer et al., 2019), so that the rotation of the ball measured the fly’s intended turning behavior. Full-field visual motion elicits optomotor turning behavior in flies, in which they turn in the direction of perceived motion (Borst & Egelhaaf, 1989; Egelhaaf et al., 1989; Fermi & Reichardt, 1963; Götz, 1964; Hassenstein & Reichardt, 1956; Kunze, 1961; McCann & MacGinitie, 1965; Reichardt & Poggio, 1976; Reichardt & Varju, 1959). Thus, their turn direction and speed during stimulus presentation can be interpreted as the direction and intensity of perceived visual motion. Because sinusoidal gratings are periodic, a given shift can in principle be interpreted as motion along many possible spatial paths: the shift plus or minus any whole number of wavelengths. However, only two paths involve shifts of less than one wavelength: the shorter spatial displacement (short-path) and the longer spatial displacement (long-path) in the opposite direction. When there is no interleave, humans resolve the ambiguity in the displacement in favor of perceiving motion in the short-path direction, while substantial interleaves lead to percepts in the long-path direction, an effect that has been described as an illusory reversal in direction (Pantle & Turano, 1992).

To determine how the stimulus parameters affect fly behavior, we swept both the interleave durations and the phase shifts in the interleave-shift stimulus, while presenting each stationary sinusoid frame for 200ms. These stimuli were presented to flies while we recorded their turning behavior (**Fig. 1D**). We averaged the turning responses during the stimulus presentation and found that flies consistently turned in the short-path direction when presented with stimuli with short interleaves. However, as the interleave duration increased, the flies exhibited progressively less turning toward the short-path direction. For longer interleaves (e.g., 8/60 s), average turning responses were toward the long-path direction for many phase shifts (**Fig. 1E**).

These optomotor responses indicate that, similar to mammals, fly motion percepts reversed when interleaves were presented between shifts in a sinusoidal grating. Moreover, also parallel to mammalian percepts, the strength of this reversal depended on interleave duration, with perceptual reversals occurring at longer interleaves.

### The illusory motion reversal is mediated by T4 and T5 neurons

We next asked which visual circuits were required for behavioral responses to the interleave-shift stimulus. We hypothesized that it was mediated by the primary motion-detection circuits in the fly visual system, T4 and T5 (Maisak et al., 2013). The T4 and T5 neurons are the earliest direction-selective neurons in the motion detection circuit (**Fig. 2A**), and are required for optomotor rotations (Bahl et al., 2013; Maisak et al., 2013), regulating walking speed (Creamer et al., 2018), estimating visual distance (Shomar et al., 2025), and directing landing responses (Schilling & Borst, 2015).

**Figure 2.**
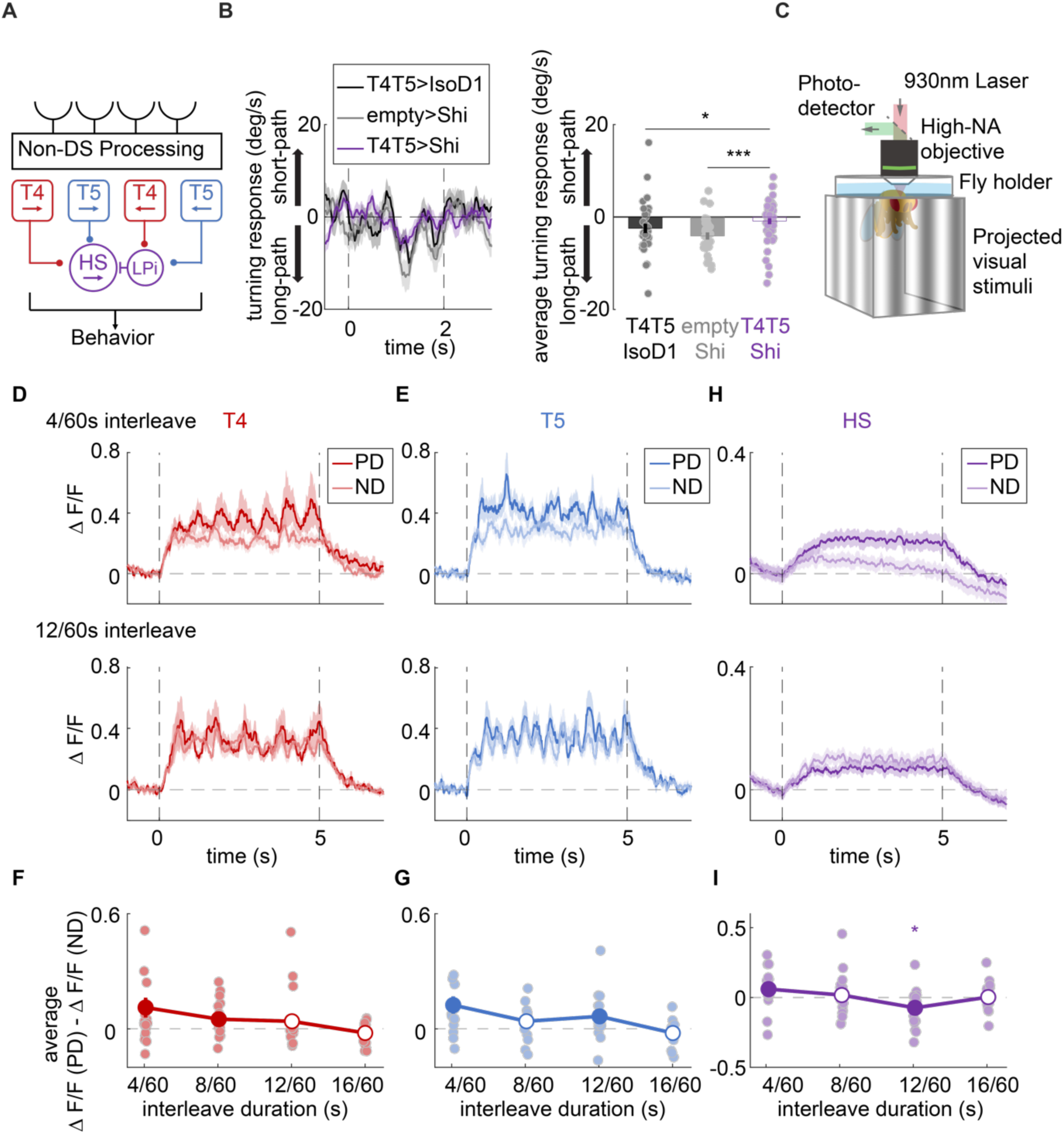
The illusory motion reversal induced by interleave-shift stimuli is mediated by T4 and T5 neurons. (A) Circuit diagram of the direction selective neurons in fly visual system. (B) Time traces (left) and averaged turning response (right) of T4-T5-silenced flies and respective control flies towards the stimuli with 7/16 cycle shift and 8/60 s interleave. (N = 43 flies for T4T5 > IsoD1; N = 33 flies for empty splitGal4 > Shi; N = 56 flies for T4T5 > Shi). Error bars indicate mean ± SEM and the dot is filled if p < 0.05 by Wilcoxon signed-rank tests against 0. * p < 0.05 and *** p < 0.001 by Wilcoxon rank-sum tests across flies. (C) Imaging setup for measuring neural responses from direction selective neurons. NA, numerical aperture. (D-E) Time traces of GCaMP6f calcium response in T4 (D) and T5 (E) neurons responding to interleave-shift stimuli shifting towards the preferred direction (PD) and null direction (ND) of neurons with 4/60 s interleave (top) and 12/60s interleave (bottom), the shifts for all stimuli are 3/8 cycle. (F-G) subtraction of averaged calcium response in T4 (G) and T5 (H) neurons towards PD-shifted stimuli and ND-shifted stimuli. Each dot represents 1 fly (N = 21 flies for T4; N = 17 flies for T5). Error bars indicate mean ± SEM and the dot is filled if p < 0.05 by Wilcoxon signed-rank tests against 0. * p < 0.05 by Wilcoxon signed-rank tests against 0, shown only for negative significant values. (H) Time traces of GCaMP7b calcium response in HS neurons responding to interleave-shift stimuli shifting towards the preferred direction (PD) and null direction (ND) of neurons with short 4/60 s interleave (top) and 12/60 s interleave (bottom), the shifts for all conditions are 3/8 cycle. (I) subtraction of averaged calcium response in HS neurons towards PD-shifted stimuli and ND-shifted stimuli. Each dot represents 1 fly (N = 19 flies). Error bars indicate mean ± SEM and the dot is filled if p < 0.05 by Wilcoxon signed-rank tests against 0.

To test this hypothesis, we examined how silencing T4 and T5 neurons affects the illusory motion reversal induced by the interleave-shift stimulus. We silenced T4 and T5 neurons using Shibire^ts^, a temperature-sensitive suppressor of synaptic transmission (Kitamoto, 2001) while presenting the interleave-shift stimulus. Silencing these neurons abolished the turning responses to the stimulus in the long-path direction (**Fig. 2B**), consistent with the broad role of T4 and T5 in motion-guided behaviors. This demonstrates that the perceptual reversal of motion in the interleave-shift stimulus depends critically on the activity of T4 and T5 neurons.

To understand how these neurons mediate this response, we measured their responses to the stimulus. We expressed the calcium indicator GCaMP6f (Chen et al., 2013) in T4 and T5 neurons, then used a two-photon microscope to record the fluorescence in these neurons while presenting the fly with interleave-shift stimuli (**Fig. 2C**). As in the behavioral assays with strong turning responses in the long-path direction, the stimuli were sinusoids with 30° wavelength and a 3/8-cycle shift, with varied interleave durations and shift directions.

T4 and T5 neurons respond strongly to widefield motion in their preferred direction (PD) and little to motion in the opposite, null direction (ND) (Maisak et al., 2013). We examined the responses of T4 and T5 to interleave-shift stimuli with shifts oriented in their PD and ND. To do so, we defined the shift direction as the direction of the shortest phase shift across interleaves. Stimuli shifting in both the PD and ND elicited robust calcium responses in T4 and T5 neurons (**Fig. 2D-E**), consistent with prior measurements of T4 and T5 responses to flickering sinusoidal stimuli (Badwan et al., 2019; Fisher et al., 2015; Wienecke et al., 2018). These results suggest that these motion detectors are sensitive to both the long-path and short-path directions of motion.

We therefore compared the time-averaged calcium responses evoked by stimuli shifting toward the PD and the ND. With short interleaves (e.g., 4/60s), responses to the PD-shifted stimuli were consistently stronger than those to the ND. However, as the interleave duration increased, this difference diminished (**Fig. 2F-G**), indicating that longer interleaves effectively decrease direction-selectivity of T4 and T5 neurons to interleave-shift stimuli.

The calcium response of these motion detectors to the interleave-shift stimuli is similar to the results of a simulation of the motion energy model (**Fig. S1**) (Adelson & Bergen, 1985), which has been proposed to account for this perceptual reversal (Strout et al., 1994; Takeuchi & De Valois, 1997). This simulation of the motion energy model used spatiotemporal filters designed to mirror the measured ones (Arenz et al., 2017; Gruntman et al., 2018, 2019; Leong et al., 2016; Salazar-Gatzimas et al., 2016) (see **Methods**). This simple energy model applied to reasonable spatiotemporal filters could result in the reversal of direction with long interleaves for interleave-shift sinusoidal stimuli. In this model, the steady state response of the motion detectors to a stimulus is a weighted sum of the stimulus motion energy across spatial and temporal frequencies, with the weighting determined by the spatiotemporal filter shape. Without interleaves, short-path motion typically dominates because of its higher energy and greater receptive field weight (**Fig. S1K and M**). Introducing longer interleaves reduces the temporal frequencies of all motion components, which increases the receptive field weight for long-path motion relative to short-path motion (Fig. **S1L-M)**. This shift results in higher weighted motion energy for the long-path motion, thereby producing motion reversal (**Fig. S1M**).

Our observation that T4 and T5 neurons respond to both PD and ND stimuli, with diminished difference at longer interleaves, is consistent with the simulation where the interleave-shift stimulus activates motion detectors for both directions (**Fig. S1G-L**) and that their relative activity shifts with interleave duration (**Fig. S1M**). However, the simulation predicted that long-path motion detectors should become dominant at longer interleaves, a pattern not clearly reflected in T4 or T5 calcium signals. This discrepancy may stem from potential differences between measured calcium signals and outputs to downstream neurons, for instance caused by synaptic release that scales super-linearly with calcium concentration (Dodge & Rahamimoff, 1967). Indeed, applying a simple quadratic transformation to T4 and T5 time traces before averaging could generate a net reversal at long interleaves (**Fig. S2**), supporting this possibility.

To further test whether the motion reversal arises from integrating signals from different direction selective T4 and T5 neurons, we recorded activity from a primary downstream target of T4 and T5, the horizontal system (HS) neurons (Hausen, 1982a, 1982b), which receive direct excitatory input from T4 and T5 with one directional tuning and indirect inhibitory input from T4 and T5 with the opposite directional tuning (Mauss et al., 2015; Schnell et al., 2010). The HS neurons are selective for horizontal motion directions and their activity is sufficient to elicit turning (Haikala et al., 2013). Using GCaMP7b (Dana et al., 2019), we measured calcium responses in HS neurons, which are depolarized by PD motion and hyperpolarized by ND motion (Hausen, 1982a, 1982b). GCaMP7b has a lower K_d_ than GCaMP6f (Dana et al., 2019), potentially permitting observations of neural hyperpolarization in its calcium signals.

When presenting the same stimuli as we presented to T4 and T5, we found that with short interleaves (e.g., 4/60 s), PD-shifted stimuli elicited stronger calcium responses in HS neurons than ND-shifted stimuli. Conversely, with long interleaves (e.g., 12/60 s), ND-shifted stimuli evoked significantly stronger responses in HS neurons than PD-shifted stimuli (**Fig. 2F and I)**, in qualitative agreement with the behavioral results (**Fig. 1**). The tuning in behavior was to shorter interleaves than elicited reversals in imaging, a difference that may be related to tuning changes that depend on fly behavior (Arenz et al., 2017; Chiappe et al., 2010; Strother et al., 2018). Overall, the reversal in HS neuron signals reflects the biological integration of motion signals from T4 and T5, which may be more sensitive than our artificial subtraction of calcium signals from T4 and T5 cells with opposite direction tuning (**Fig. 2F-G and I**). These results in HS are also consistent with the motion energy account of the perceptual reversal (**Fig. S1**).

### After images intuitively account for direction reversals

In parallel to the motion energy explanation, a non-exclusive account based on afterimages has been proposed to explain the reversal of mammalian motion percepts evoked by the interleave-shift stimuli (Miura et al., 2018; Ohnishi et al., 2016; Sheliga et al., 2006; Sugita et al., 2020). Afterimages occur after prolonged exposure to an image; when the image is removed, visual systems generate a contrast-inverted afterimage of the original. In this account of the perceptual reversal, in the interleave-shift stimuli, when sinusoidal gratings are replaced by a uniform, mean-luminance interleave, animals see an inverted afterimage of the gratings during the interleave (**Fig. 3A**). The shortest path direction between the inverted afterimage during the interleave and the subsequent shifted frame is in the direction of the long path between the two sinusoidal frames (**Fig. 3A**). If observers see motion in the short-path direction between afterimage and subsequent frames, this can account for the illusory direction reversal. In flies, afterimages are known to influence neural responses to motion (Egelhaaf & Borst, 1989; Harris & O’Carroll, 2002).

**Figure 3.**
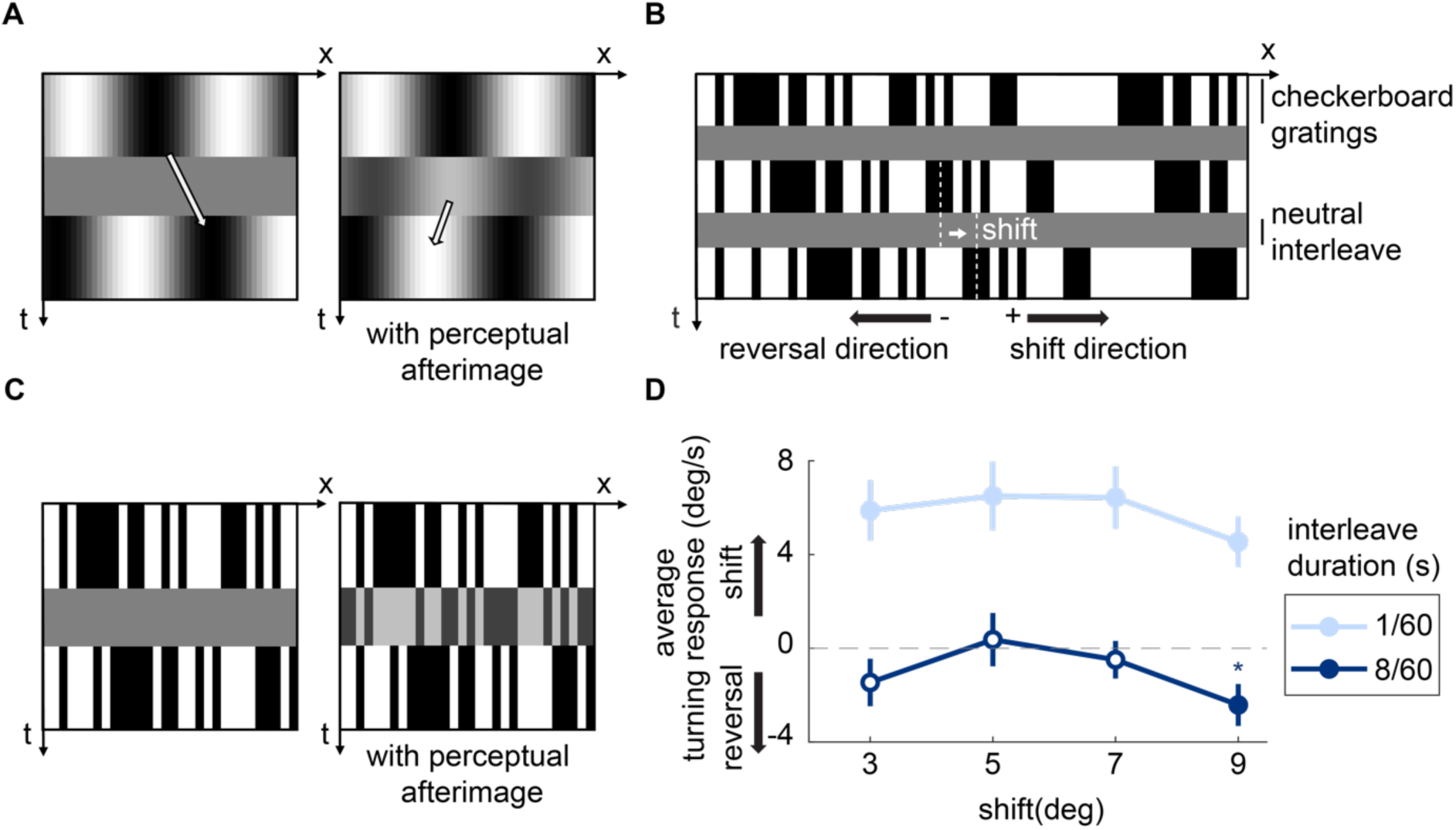
The illusory motion reversal induced by interleave-shift stimulus can be explained by afterimages. (A) Space-time contrast diagram of the interleave-shift stimulus (left) and an illustration of the inverted afterimage that may exist during interleaves (right). The arrows indicate the shortest path that the sinusoid gratings move. (B) Diagram of the checkerboard interleave-shift stimulus that is adapted from the original sinusoid interleave-shift stimulus. (C) Diagram of the checkerboard interleave-shift stimulus (left) and an illustration of the proposed inverted afterimage perceived in interleaves (right). (D) Averaged turning response of wild-type flies towards checkerboard interleave-shift stimuli with different shifts and interleave durations (N = 44 flies). Error bars indicate mean ± SEM and the dot is filled if p < 0.05 by Wilcoxon signed-rank tests against 0. * p < 0.05 by Wilcoxon signed-rank tests against 0, shown only for negative significant values.

To test this hypothesis, we created a new interleave-shift stimulus in which the periodic sinusoidal gratings were replaced by random binary gratings (**Fig. 3B**). We hypothesized that this stimulus would elicit a similar direction reversal effect, despite the lack of spatial periodicity. According to the afterimage explanation, after prolonged presentation of the checkerboard pattern, replacing it with a uniform interleave should result in an inverted afterimage (**Fig. 3C**). This inverted afterimage, being anticorrelated with the subsequent shifted checkerboard pattern, could then induce a *reverse-phi* illusory motion percept (Hassenstein & Reichardt, 1956). This reverse-phi phenomenon refers to the perception of motion in the opposite direction when an image is shifted and inverted in polarity (Anstis, 1970; Hassenstein & Reichardt, 1956). This phenomenon has been widely used to probe motion detection algorithms and mechanisms (Adelson & Bergen, 1985; Anstis, 1970; Bours et al., 2009; Hassenstein & Reichardt, 1956; Orger et al., 2000; Salazar-Gatzimas et al., 2018; Salazar-Gatzimas et al., 2016; Tuthill et al., 2011; Tuthill et al., 2013). Thus, the interaction between the shifted image and the inverted afterimage could still produce motion percepts in the direction opposite the shift via the reverse-phi mechanism, but only if the interleave is long enough to generate the afterimage.

We presented these random binary grating interleave-shift stimuli with varying shifts and interleave durations in the same optomotor behavioral paradigm. At short interleaves, flies turned in the direction of the displacement (**Fig. 3D**). At a longer interleave of 8/60 s, flies turned in the direction opposite the shift at the longest shift, likely reflecting a balance between responses to the positive correlations between subsequent grating frames and negative correlations with afterimages, which could be tuned to different lengthscales or be influenced by the shape of spatial receptive fields (Arenz et al., 2017). These results mirror human studies, showing direction reversals with interleaved random binary gratings (Nohara et al., 2015) and dots (Shioiri & Cavanagh, 1990). This finding demonstrates that non-periodic interleaved checkerboard stimuli can evoke the illusory motion reversal, consistent with the afterimage account for the illusory reversal.

### Afterimages are widespread in the motion-detection neural circuit

To further investigate how afterimages arise in the fly’s visual circuits, we measured from neurons while presenting flies with a stimulus consisting of a stationary sinusoidal grating followed by a uniform neutral contrast (**Fig. 4A**). (We refer to the interleave as neutral if it is mean-luminance; the Weber contrast of the interleave is computed relative to the mean of the sinusoidal frames.) This pattern-uniform stimulus mimicked the transition from sinusoidal gratings to interleaves, where afterimages are likely to occur. We focused on neurons upstream of T4 and T5, which process contrast information using distinct temporal filters before transmitting signals to T4 and T5. Specifically, we examined Mi1 and Tm3 (ON cells upstream of T4 neurons) and Tm1 and Tm2 (OFF cells upstream of T5 neurons) (**Fig. 4B**) (Shinomiya et al., 2019; Strother et al., 2017; Takemura et al., 2017) , as previous studies have shown these neurons employ derivative-taking temporal filters (Arenz et al., 2017; Behnia et al., 2014; Strother et al., 2017).

**Figure 4.**
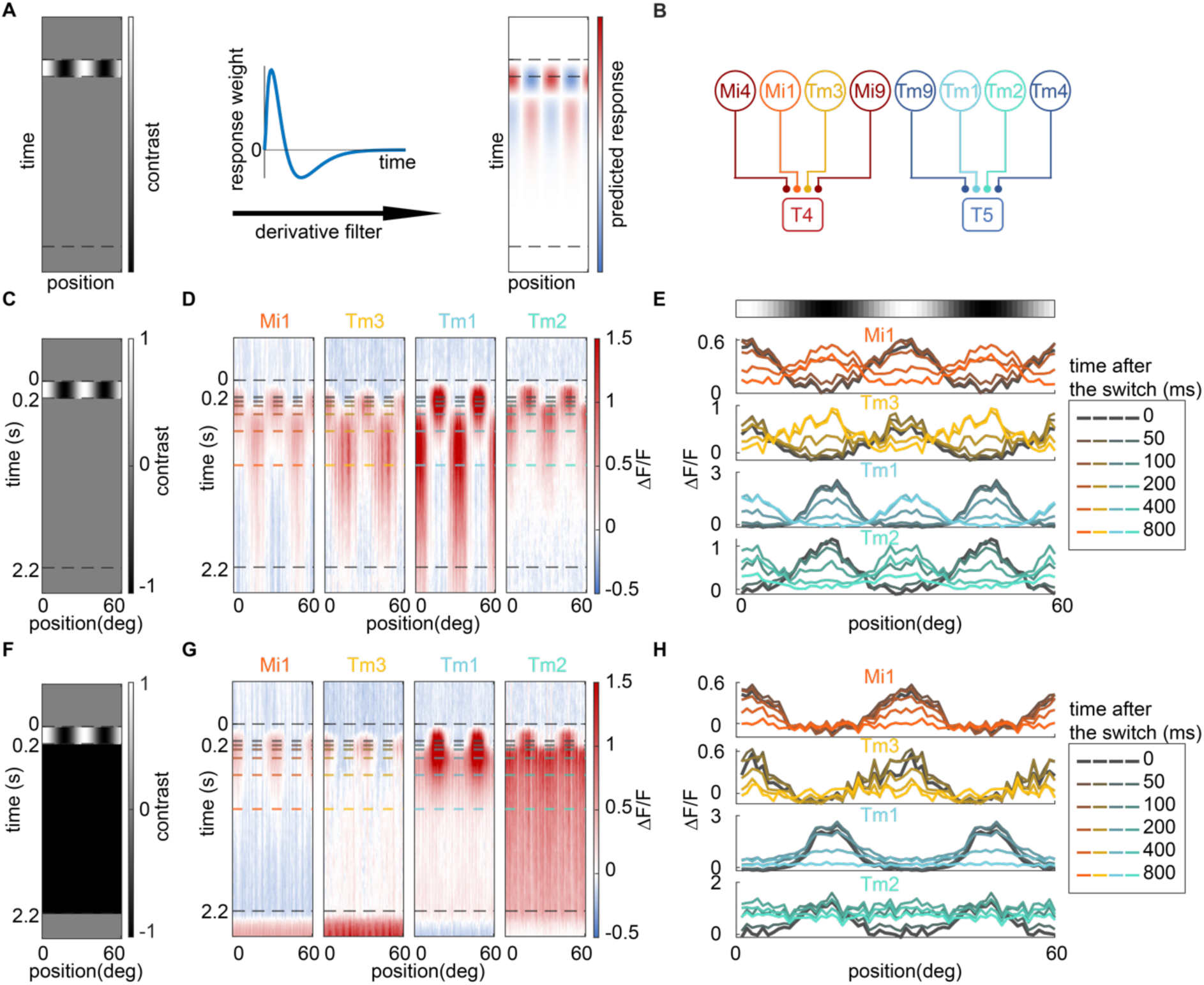
Afterimages are widely present in the neurons upstream of the direction selective circuit. (A) Space-time contrast diagram of a sinusoidal pattern-uniform stimulus switching from sinusoid gratings to a uniform neutral interleave (left) and the simulated neural response towards it (right), generated by filtering the stimulus by a temporal derivative-taking temporal filter. (B) Circuit diagram of upstream neurons that process contrast information using distinct linear temporal filters before transmitting signals to T4 and T5. Synapses are denoted by dots regardless of synapse sign. (C-D) Diagram of a sinusoidal pattern-uniform stimulus switching to neutral interleave (C) and calcium responses of Mi1, Tm3, Tm1, Tm2 (D). For each neuron type, the ROIs are assigned to a phase by mapping their receptive field (see Methods). The ROIs are aligned based on their phases, the heatmaps are replicated 2 cycles for better visualization. (E) Calcium responses of Mi1, Tm3, Tm1, Tm2 at different times after the sinusoid gratings switch to neutral interleave. (F-G) Diagram of a sinusoidal pattern-uniform stimulus switching to a dark interleave (F) and calcium responses of Mi1, Tm3, Tm1, Tm2 (G). (H) Calcium response of Mi1, Tm3, Tm1, Tm2 at different times after the sinusoid gratings switch to the dark interleave. (n = 88 ROIs from 4 flies for Mi1; n = 69 ROIs from 2 flies for Tm3; n = 106 ROIs from 4 flies for Tm1; n = 79 ROIs from 6 flies for Tm2)

Using GCaMP6f to record calcium activity, we aligned the regions of interest by their azimuthal receptive field position. During grating presentation, all four neuron types exhibited spatial activity patterns consistent with their ON and OFF sensitivity. But upon transition to the uniform neutral interleave (**Fig. 4C**), these neurons showed inverted response patterns, or afterimages (**Fig. 4D-E**).

To test whether this afterimage-like response is driven by derivative-taking filters, we conducted the same experiment on Mi4 and Mi9 neurons, which are characterized as integrating signals over time rather than taking derivatives (Arenz et al., 2017). In contrast to the other four neurons, these neurons did not exhibit strong afterimage-like response patterns (**Fig. S3B-C**), indicating that afterimages do not occur in every neuron type upstream of T4 and T5.

The inverted afterimage-like pattern did not appear immediately following the contrast transition, but rather after a delay of ∼0.4 seconds (**Fig. 4E**). The measured delay in these experiments likely depends on both cellular processing (Arenz et al., 2017; Gonzalez-Suarez et al., 2022; Yang et al., 2016) and sensor dynamics (Chen et al., 2013), but the delayed emergence of afterimage responses also explains why longer interleaves lead to stronger long-path motion percepts in response to the sinusoidal interleave-shift stimulus.

### Suppressing afterimages by using dark interleaves eliminates the illusory motion reversal

We discovered that the afterimages observed in neurons upstream of T4 and T5 depended critically on the stimulus contrast. When we presented upstream neurons with the sinusoidal pattern-uniform stimuli that transitioned to a dark uniform interleave (**Fig. 4F**), they did not display afterimage-like responses (**Fig. 4G-H**), even though they had afterimage-like responses when the interleave was neutral. Conversely, when the transition was to a light uniform interleave (**Fig. S3E**), these neurons showed weak afterimage-like responses (**Fig. S3F-G**). These findings indicate that the contrast of the pattern-uniform stimulus strongly influences the magnitude of the afterimage effect, with dark interleaves effectively abolishing it.

This observation led us to hypothesize that perception of the interleave-shift stimulus could be manipulated by adjusting the contrast of the interleaves. If the illusion indeed depends on the presence of afterimages, then replacing neutral interleaves with dark ones should suppress the motion reversal. In this case, even with long interleave durations, the perceived motion should remain in the short-path direction, since the afterimages would be eliminated from the direction selective circuit.

To test this hypothesis, we conducted both neural imaging and behavioral experiments using modified stimuli in which the interleaves were dark (–100% contrast) instead of neutral. At the neuronal level, we presented dark interleaves between shifted sinusoids while measuring from HS neurons (**Fig. 5A-B**). Under these conditions, HS neurons did not show the reversal they did to neutral interleaves. Thus, the reversed neural response observed in the original interleave-shift stimulus with neutral interleaves was abolished when the afterimage effect is eliminated by employing dark interleaves.

**Figure 5.**
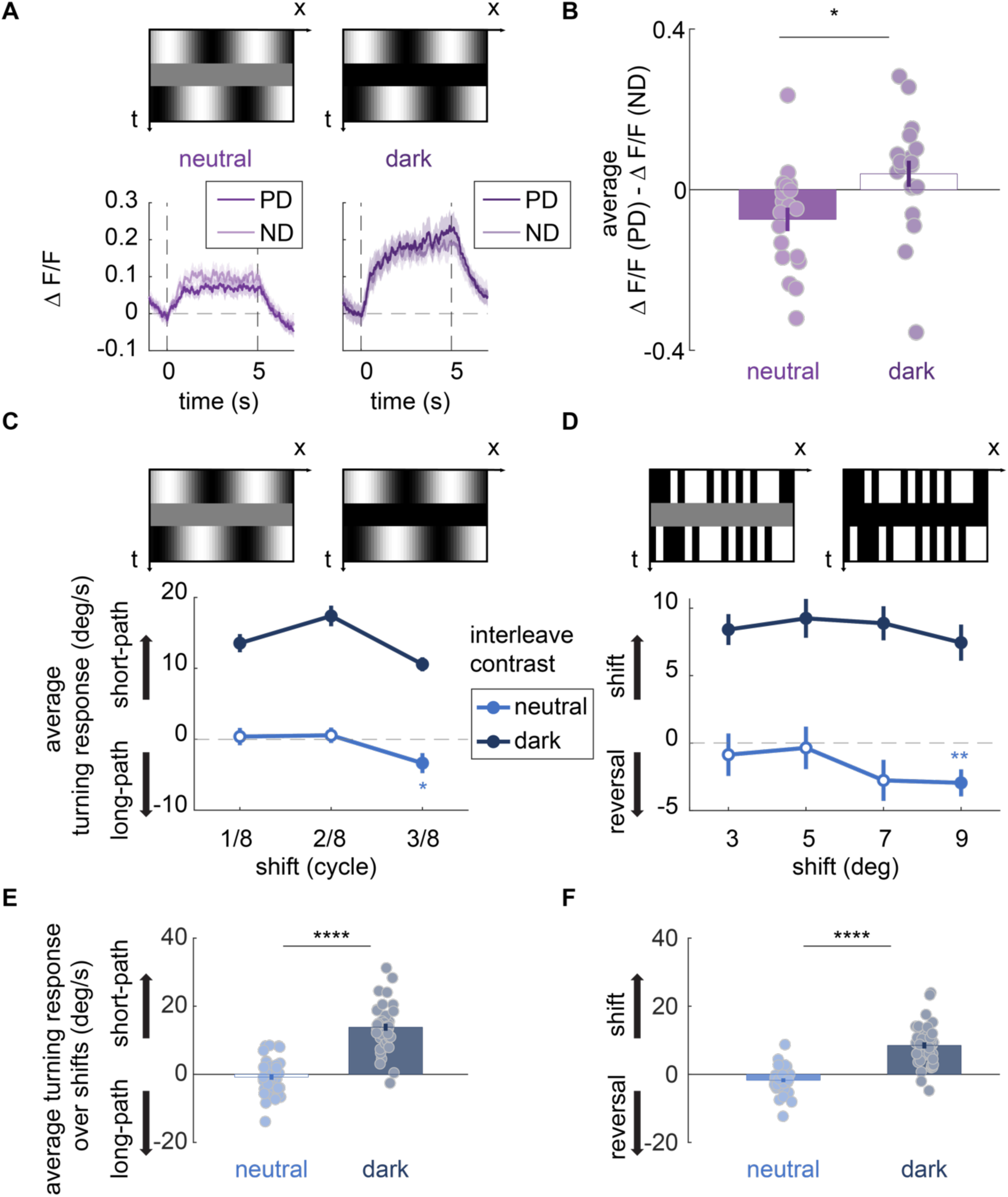
Reducing afterimages using dark interleaves decreases the illusory motion reversal. (A) Time traces of calcium response in HS neurons responding to PD-shifted and ND-shifted interleave-shift stimuli with neutral (left) and dark (right) 12/60 s interleave. The shifts for all stimuli ere 3/8 cycle. (B) Difference of averaged calcium response in HS neurons to PD-shifted stimuli and ND-shifted stimuli with neutral and dark interleaves (data for neutral interleave condition is the same from Fig. 2H**-I**). Each light dot represents 1 fly (N = 19 flies). Error bars indicate mean ± SEM and the box is filled if p < 0.05 by Wilcoxon signed-rank tests against 0. * p < 0.05 by Wilcoxon signed-rank tests across interleave contrast conditions. (C-D) Averaged turning response of wild-type flies towards sinusoid (C) and checkerboard interleave-shift stimuli (D), both with 8/60s interleave duration but with different shifts and interleave contrasts (N = 41 flies). Error bars indicate mean ± SEM and the dot is filled if p < 0.05 by Wilcoxon signed-rank tests against 0. ** p < 0.01 by Wilcoxon signed-rank tests against 0, shown for negative significant values. (E-F) Average turning response of wild-type flies evoked by sinusoid (E) and checkerboard (F) interleave-shift stimuli over different shifts. (N = 41 flies) Error bars indicate mean ± SEM and the box is filled if p < 0.05 by Wilcoxon signed-rank tests against 0. **** p < 0.0001 by Wilcoxon signed-rank tests across interleave contrast conditions.

At the behavioral level, flies were presented with both the original interleave-shift and checkerboard interleave-shift stimuli, as well as altered versions of each that employed dark interleaves (–100% contrast) (**Fig. 5C-F**). In both cases, flies did not exhibit the reversed turning associated with the neutral interleave version of these stimuli, instead turning relatively strongly in the direction of the short-direction displacement for all interleave durations. Thus, the illusory motion reversal in both stimuli was eliminated when afterimages were suppressed by dark interleaves.

When we repeated the same experiments using light interleaves (+100% contrast), which preserve a weak afterimage (**Fig. S3**), HS responses showed attenuation of directional signals with longer interleave durations, but did not show the significant reversal that we observed with neutral interleaves (**Fig. S4A-B**). Interestingly, at the behavioral level, flies exhibited substantially stronger reversals with light interleaves compared to neutral ones (**Fig. S4C-F**). This discrepancy suggests the presence of an unidentified pathway that enhances behavioral output under the light interleave condition.

Together, these findings demonstrate that eliminating afterimages by using dark interleaves abolishes the illusory motion reversal, both at the neural and behavioral levels. Conversely, preserving afterimages with light interleaves maintains or enhances the reversal.

## Discussion

These results demonstrate that, like mammals, fruit flies perceive an illusory motion reversal induced by interleave-shift stimuli. Moreover, in flies as in mammals, the strength of this illusion increases with the interleave duration. This illusion joins two other motion illusions shared by insects and humans: reverse-phi percepts (Anstis, 1970; Hassenstein & Reichardt, 1956) and peripheral drift illusions (Agrochao et al., 2020; Fraser & Wilcox, 1979). The reverse-phi illusion highlights how pairwise correlations are used to generate motion signals, consistent with correlation detectors and the motion energy model (Adelson & Bergen, 1985; Hassenstein & Reichardt, 1956). In contrast, peripheral drift illusions rely on light-dark asymmetries and therefore cannot be explained by canonical correlation or energy models (Agrochao et al., 2020; Faubert & Herbert, 1999; Fitzgerald et al., 2011). Here, we showed that the direction reversal of percepts of the interleave-shift stimulus can be only partially accounted for by the motion energy model, and that light-dark asymmetries play an important role.

Interestingly, one of the interleave-shift stimuli we used highlights the interaction between afterimages and the reverse-phi illusion. When we presented and shifted stochastic binary patterns (**Fig. 3B**), the afterimages during the interleave had the inverse contrast pattern of the prior binary pattern. Reverse-phi percepts occur when a stimulus inverts its contrast when it is displaced over time — in such cases, percepts are of motion in the direction opposite the displacement (Anstis, 1970; Hassenstein & Reichardt, 1956). Thus, when these stochastic binary interleave-shift stimuli caused flies to turn in the direction opposite displacement, it can be viewed as a reverse-phi percept generated between the afterimage during the interleave and the subsequent, shifted frame. In that sense, the reversal is the result of both an afterimage and sensitivity to negative spatiotemporal correlations.

### Motion energy and afterimages offer complementary accounts for this phenomenon

The motion energy model has been proposed as an explanation of this illusion in earlier studies (Strout et al., 1994; Takeuchi & De Valois, 1997). According to this model, increasing interleave duration increases the relative weighting of long-path motion energy, leading to the perception of reversed motion (**Fig. S1**). However, it is notable that the differences in behavior between the light and dark interleaves are inconsistent with a canonical motion energy model. This is because stimuli with light and dark interleave contrasts (**Fig. S5A-C**) have identical power, or energy, in spatiotemporal frequency space (**Fig. S5D-F**). According to the motion energy model, stimuli with identical motion energy, such as the light and dark interleave versions of the stimulus, should evoke similar percepts (**Fig. S5**), yet we observed starkly different behavioral responses in the reversal (**Fig. 5C-F, Fig. S4C-F**).

These inconsistencies are better explained by the afterimage account, which does not rely on light-dark symmetry, as the motion energy model does (Clark et al., 2014; Fitzgerald et al., 2011). The afterimages explanation is not exclusive of motion energy, but if one allows afterimages to appear under some conditions but not others, then it is less rigid than the motion energy model and effectively permits different temporal processing for different interleave contrasts.

### Light-dark asymmetries in motion detection

While both flies and humans exhibit similar the illusory motion reversals to the interleave-shift stimulus, a key difference between them is the effect of interleave contrast. In humans, the illusion is reported abolished with both light and dark interleaves (Pantle & Turano, 1992). Meanwhile in flies, the illusion is abolished only by dark interleaves and strengthened by light interleaves, showing a clear light-dark asymmetry.

This asymmetry is consonant with a host of other light-dark asymmetries in fly behavioral and neural responses to motion (Chen et al., 2019; Clark et al., 2014; Leonhardt et al., 2016; Silies et al., 2013; Wu et al., 2025). However, this asymmetry does not appear be directly related to differential involvement of the T4 ON pathway and T5 OFF motion pathways. For example, we observed similar responses in T4 (ON pathway) and T5 (OFF pathway) neurons (**Fig. 2**). Likewise, we observed similar afterimage patterns in Mi1, Tm3 (ON) and Tm1, Tm2 (OFF) neurons (**Fig. 4**, **Fig. S3**), including a similar loss of afterimage effects with a dark uniform screen for both ON and OFF pathway neurons.

This loss of afterimages when switching to a dark, uniform stimulus is not explained by simple linear filtering, which would preserve afterimages under all uniform contrasts (**Fig. S3A, D**). These results suggest that dark interleaves disrupt the normal processing of contrast history in derivative-taking neurons. Similar disruptions may be observed in electrical recordings of these neurons when contrasts change to their darkest levels (Behnia et al., 2014). The darkest stimuli also recruit different early visual circuitry in the fly (Ketkar et al., 2022), so that the contrast differences observed may relate to luminance gain control processing (Ketkar et al., 2023; Ketkar et al., 2020).

### Resolving perceptual ambiguity

The sinusoidal interleave-shift stimulus (**Fig. 1A-B**) is fundamentally ambiguous for underlying grating displacements. Rather than examining processing steps, one may instead adopt a Bayesian view and suppose that animals are integrating visual evidence with priors about motion to generate their percepts and behavior (Brascamp & Shevell, 2021; Knill & Richards, 1996; Von Helmholtz, 1867). In this case, if the visual system had a prior for the slowest speeds, as has been suggested in other research (Stocker & Simoncelli, 2005), then one would expect the motion direction to always be in the short-path direction for all interleave durations, contrary to what is observed. Instead, the direction reversal is consistent with a prior that favors slow speeds consistent with the stimulus, but not arbitrarily slow speeds, selecting the long-path direction—the slightly faster choice—when there are longer interleave durations. This view is consistent with the spatiotemporal tuning of motion detection representing a sort of evolved prior for behaviorally relevant motion.

### Circuitry underlying illusory percepts

Overall, by using flies as a model system, we discovered circuit- and neuron-level mechanisms underlying the perceptual direction reversal to interleave-shift stimuli (**Fig. 2**). We identified neuronal responses that support the motion energy model (**Fig. 2, Fig. S1**), which has been widely used to account for this illusion but previously lacked direct support in neural signals (Pantle & Turano, 1992; Strout et al., 1994; Takeuchi & De Valois, 1997). Moreover, characterizing afterimages has linked afterimage-based accounts of the illusion (Miura et al., 2018; Ohnishi et al., 2016; Sheliga et al., 2006; Sugita et al., 2020) to contrast-dependent effects described in other studies (**Fig. 5C-F, Fig. S4C-F**) (Bruno et al., 2011; Pantle & Turano, 1992; Shioiri & Cavanagh, 1990). The suite of perceptual phenomena shared by flies and humans highlights algorithmic processing similarities between divergent species and the advantages of cross-species comparisons for connecting circuit processing to motion detection algorithms and percepts.

## Acknowledgments

This project was funded by NIH R01 EY026555.

## Author contributions

HW and DAC conceived the behavioral and physiological experiments. HW obtained and analyzed behavioral data and neural imaging data. HW, TG, and DAC developed simulations, which were implemented by HW and TG. HW created figures and wrote the initial draft of the paper. HW and DAC edited the paper.

---

**Figure S1.**
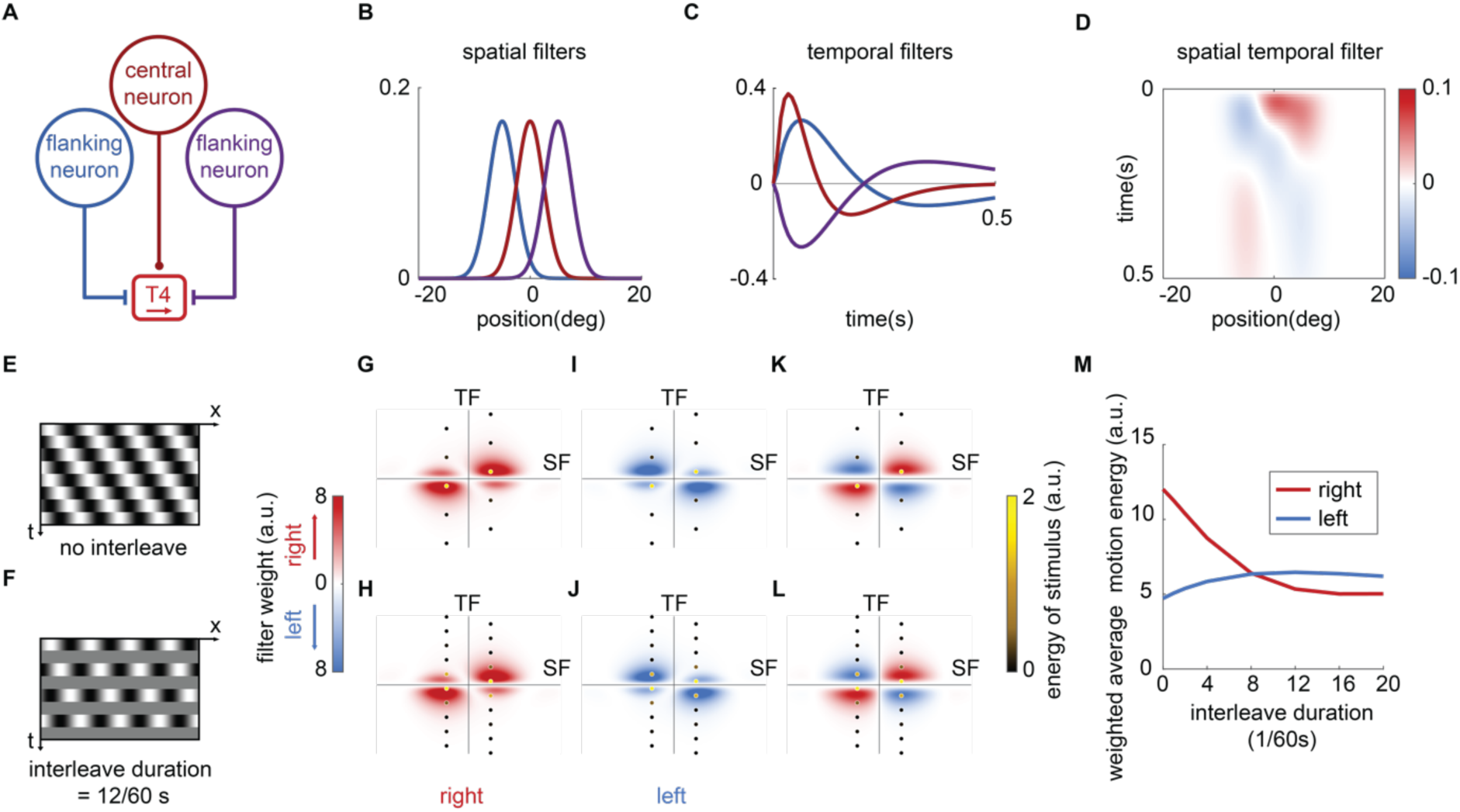
Motion energy model accounts for the interleave-motion reversal illusion with limited explanatory power. (A) Circuit diagram of a motion detecting unit in T4 neuron, which receives a positive central input signal from an upstream, ON-polarity excitatory neuron; a negative flanking input signal on the right from an upstream inhibitory, ON-polarity neuron; and a net positive flanking input signal to contrast on the left from an upstream inhibitory OFF-polarity neuron. (B) The spatial filters of upstream neurons and their relative spatial distribution. All signals share an identical Gaussian spatial filter. Each plot corresponds to the neuron in **Fig. S1A** by color scheme. (C) The temporal filters of the upstream neurons. The central signal passes through a faster temporal filter, while the side signals pass through slower temporal filters. Each plot corresponds to the neuron in **Fig. S1A** by color scheme. (D) The spatial-temporal filter of the motion detector obtained by combining the spatial and temporal filters. (E-D) Space-time diagram of shifting sinusoid gratings with 0 interleaves (E) and interleave-shift stimulus with 12/60s neutral interleaves (F). (G-L) The heatmaps represent the filter weights of rightward motion detectors (G-H), leftward motion detectors (I-J) and the combined opponent motion detectors (K-L) as a function of spatial frequency (SF) and temporal frequency (TF). Stimulus energy is overlaid as discrete points at their corresponding SF and TF coordinates for stimuli with no interleaves (G, I and K) and with 12/60 s interleaves (H, J and L). The filter weight and the energy of stimulus as a function of spatial-temporal frequency are generated by performing a Fourier transform on the space-time diagram of the filters and the stimuli (see Methods). (M) The weighted average motion energy for different directions and interleave durations.

**Figure S2.**
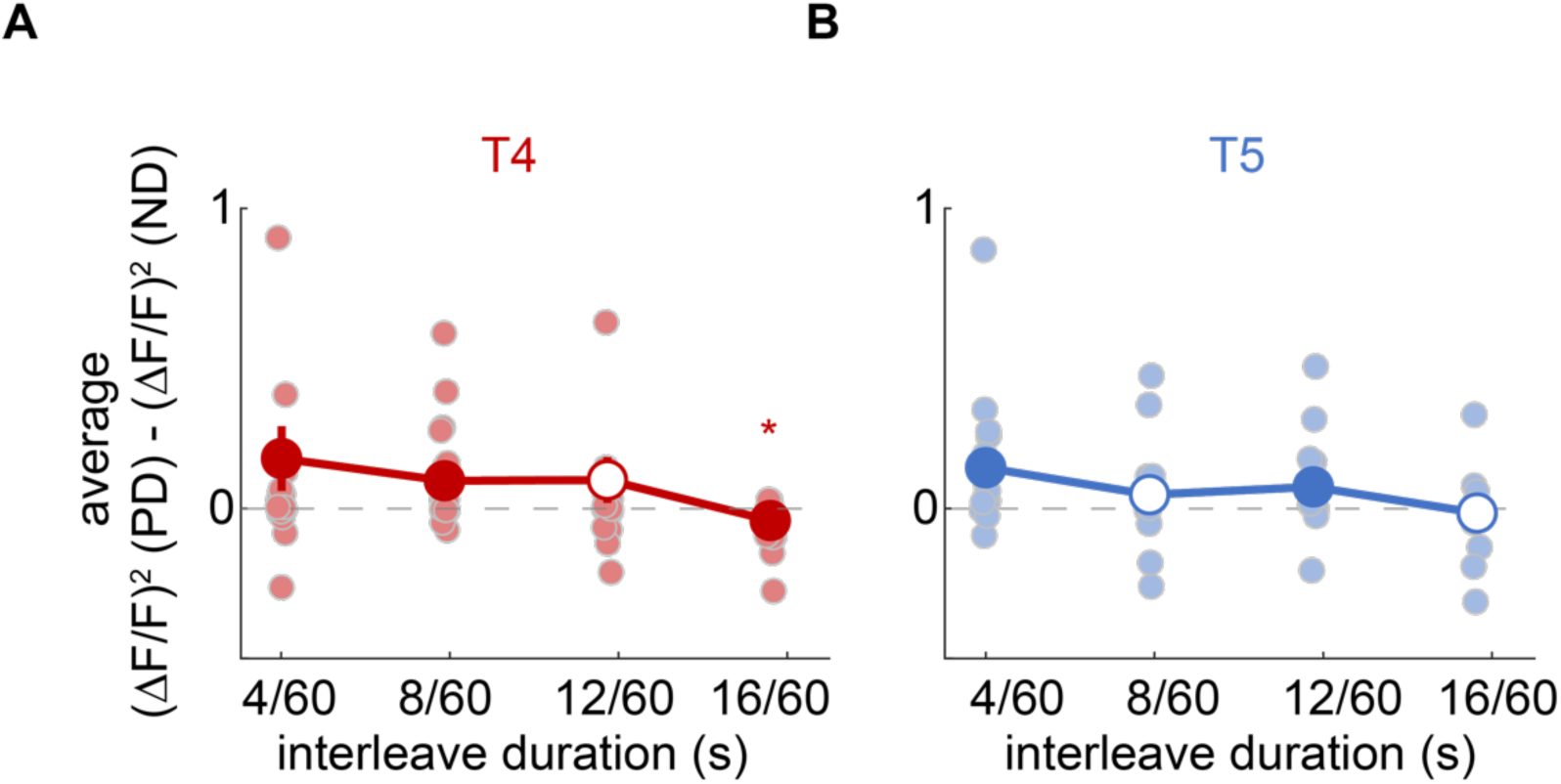
Super-linear transformations of T4T5 calcium signals before averaging could reproduce the observed reversal in relative response to PD and ND-shifted interleaved-shift stimulus. (A-B) Subtraction of averaged squared calcium response in T4 (A) and T5 (B) neurons to PD-shifted stimuli and ND-shifted stimuli. Each light dot represents 1 fly (N = 21 flies for T4; N = 17 flies for T5). Error bars indicate mean ± SEM and the dot is filled if p < 0.05 by Wilcoxon signed-rank tests against 0. * p < 0.05 by Wilcoxon signed-rank tests against 0, shown for negative significant values. Thus, transforming the calcium trace super-linearly can generate a net negative response in T4 at long interleave durations.

**Figure S3.**
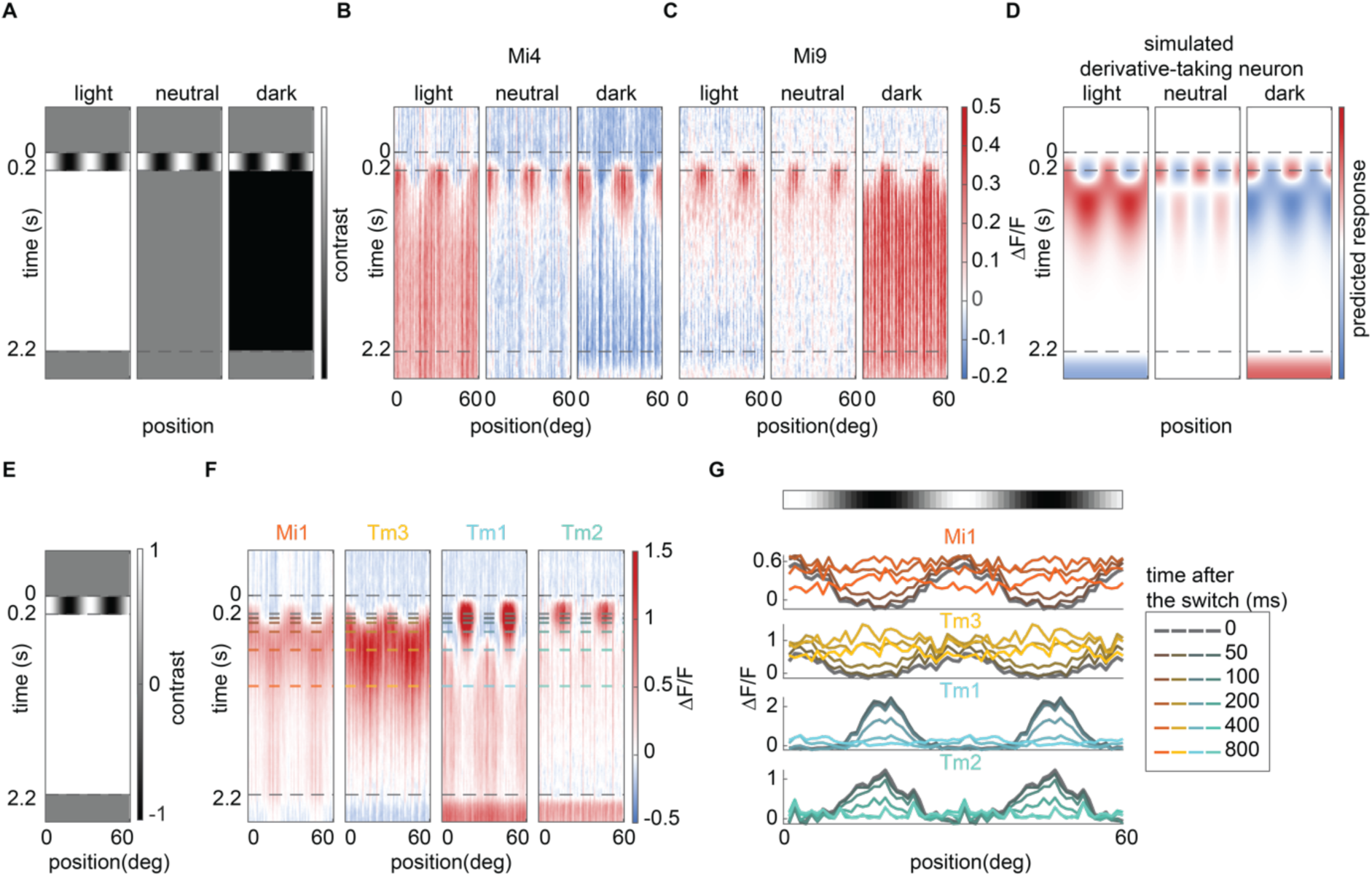
The afterimages are not present in Mi4 and Mi9 neurons and are elicited more weakly by light interleaves in neurons upstream of motion detectors. (A) Space-time contrast diagrams of sinusoidal pattern-uniform stimuli with different interleave contrasts. (B-C) Calcium responses of Mi4 (B) and Mi9 (C) evoked by sinusoidal pattern-switch stimuli with light, neutral and dark interleaves. (n = 113 ROIs from 6 flies for Mi4; n = 105 ROIs from 7 flies for Mi9) (D) Simulated responses to sinusoidal pattern-uniform stimuli with different interleave contrasts in a derivative-taking neuron, generated by filtering the stimuli with a derivative-taking temporal filter (as in Figure 4). (E, F) Diagram of a sinusoidal pattern-uniform stimulus switching to a light interleave (E) and the evoked calcium responses (F) measured in Mi1, Tm3, Tm1, Tm2. (n = 88 ROIs from 4 flies for Mi1; n = 69 ROIs from 2 flies for Tm3; n = 106 ROIs from 4 flies for Tm1; n = 79 ROIs from 6 flies for Tm2) (G) Calcium responses of Mi1, Tm3, Tm1, Tm2 at different times after sinusoid gratings switch to light interleave.

**Figure S4.**
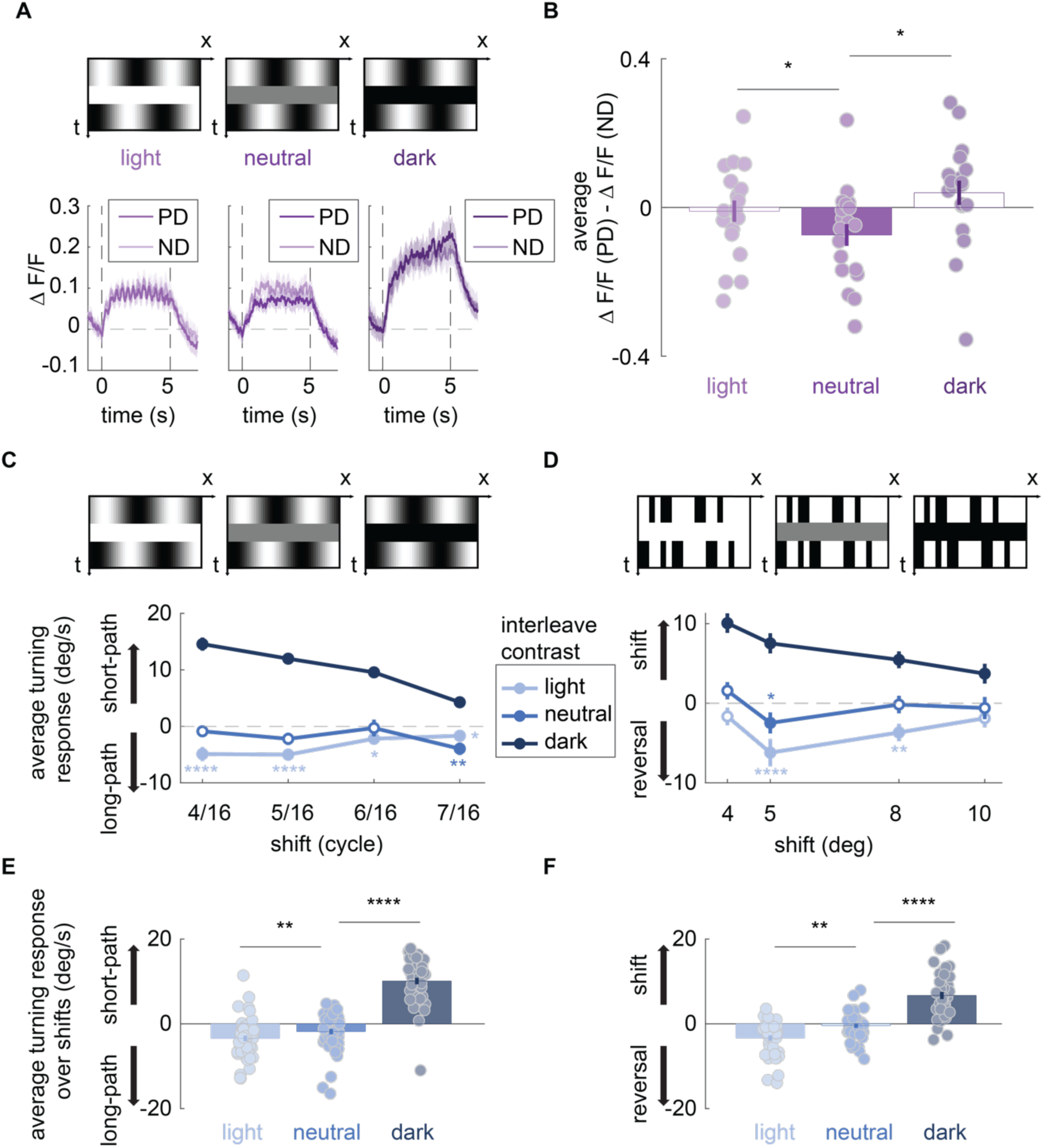
Light interleaves weaken the illusory neural response reversal in direction selective neurons, but enhance the behavioral reversal induced by interleave-shift stimulus. Time traces of calcium response in HS neurons responding to PD-shifted and ND-shifted interleave-shift stimuli with light (left), neutral (middle) and dark (right) 12/60 s interleave. The shifts for all stimuli ere 3/8 cycle. (B) Difference of averaged calcium response in HS neurons to PD-shifted stimuli and ND-shifted stimuli with light, neutral and dark interleaves (data for neutral interleave condition is the same from Fig. 2H**-I** and Fig. 5A**-B**, data for dark interleave condition is the same from Fig. 5A**-B**). Each light dot represents 1 fly (N = 19 flies). Error bars indicate mean ± SEM and the box is filled if p < 0.05 by Wilcoxon signed-rank tests against 0. * p < 0.05 by Wilcoxon signed-rank tests across interleave contrast conditions. (C-D) Averaged turning responses of wild-type flies towards sinusoid (G) and the checkerboard interleave-shift stimulus (H), both with 8/60s duration interleaves but with different shifts and interleave contrasts. (N = 50 flies for sinusoid interleave-shift stimulus; N = 41 flies for checkerboard interleave-shift stimulus) Error bars indicate mean ± SEM and the dot is filled if p < 0.05 by Wilcoxon signed-rank tests against 0. * p < 0.05, ** p < 0.01 and **** p < 0.0001 by Wilcoxon signed-rank tests against 0, shown for negative significant values. (E-F) Average turning response of wild-type flies evoked by sinusoid (I) and checkerboard interleave-shift stimuli (J) over different shifts. (N = 50 flies for sinusoid interleave-shift stimulus; N = 41 flies for checkerboard interleave-shift stimulus) Error bars indicate mean ± SEM and the box is filled if p < 0.05 by Wilcoxon signed-rank tests against 0. ** p < 0.01 and **** p < 0.0001 by Wilcoxon signed-rank tests across interleave contrast conditions.

**Figure S5.**
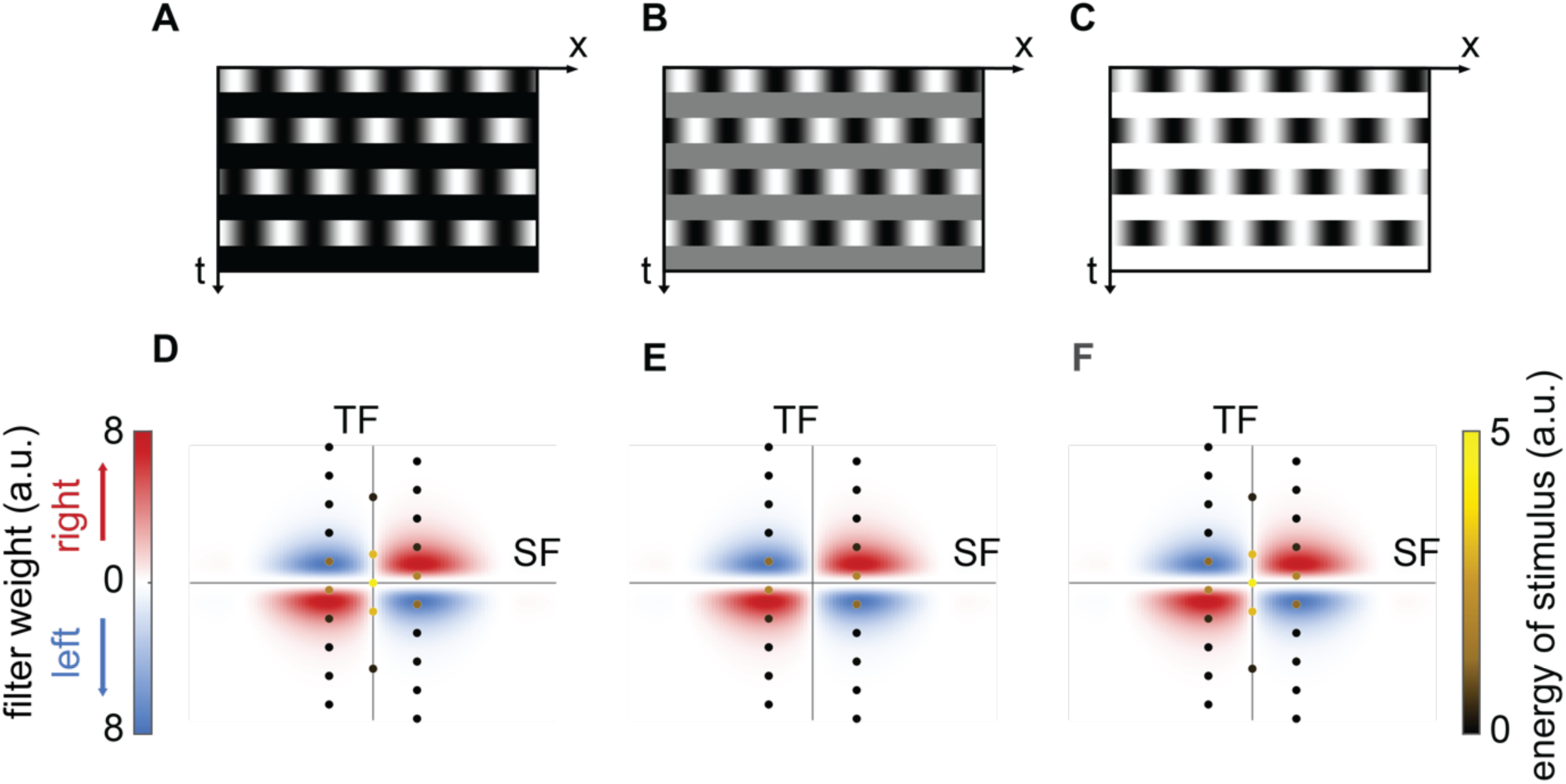
The motion energy model fails to account for the effect of interleave contrast on illusory percepts of sinusoidal interleave-shift stimuli. (A-C) Space-time diagram of interleave-shift stimuli with 12/60s dark (A), neutral (B), and light (C) interleaves. (D-F) Filter weights of the motion energy model (as in **Fig. S1**) as a function of spatial frequency (SF) and temporal frequency (TF). Stimulus energies are overlaid as points at their corresponding SF and TF coordinates for stimuli with 12/60s duration dark (D), neutral (F) and light (F) interleaves. Note that the energy in dark and light stimuli is identical.

## Methods

### Fly husbandry

Same as previous study, flies were reared at around 50% humidity on a Glucose-based food (Archon Scientific, recipe: D20102) at 20 °C for all behavioral experiments and at 25 °C for imaging experiments. Flies were reared in incubators with a 12-hour light-dark circadian cycles. Non-virgin female flies were collected using CO_2_ anesthetization within 12 hours of eclosion and were used more than 12 hours later for imaging or behavioral experiments.

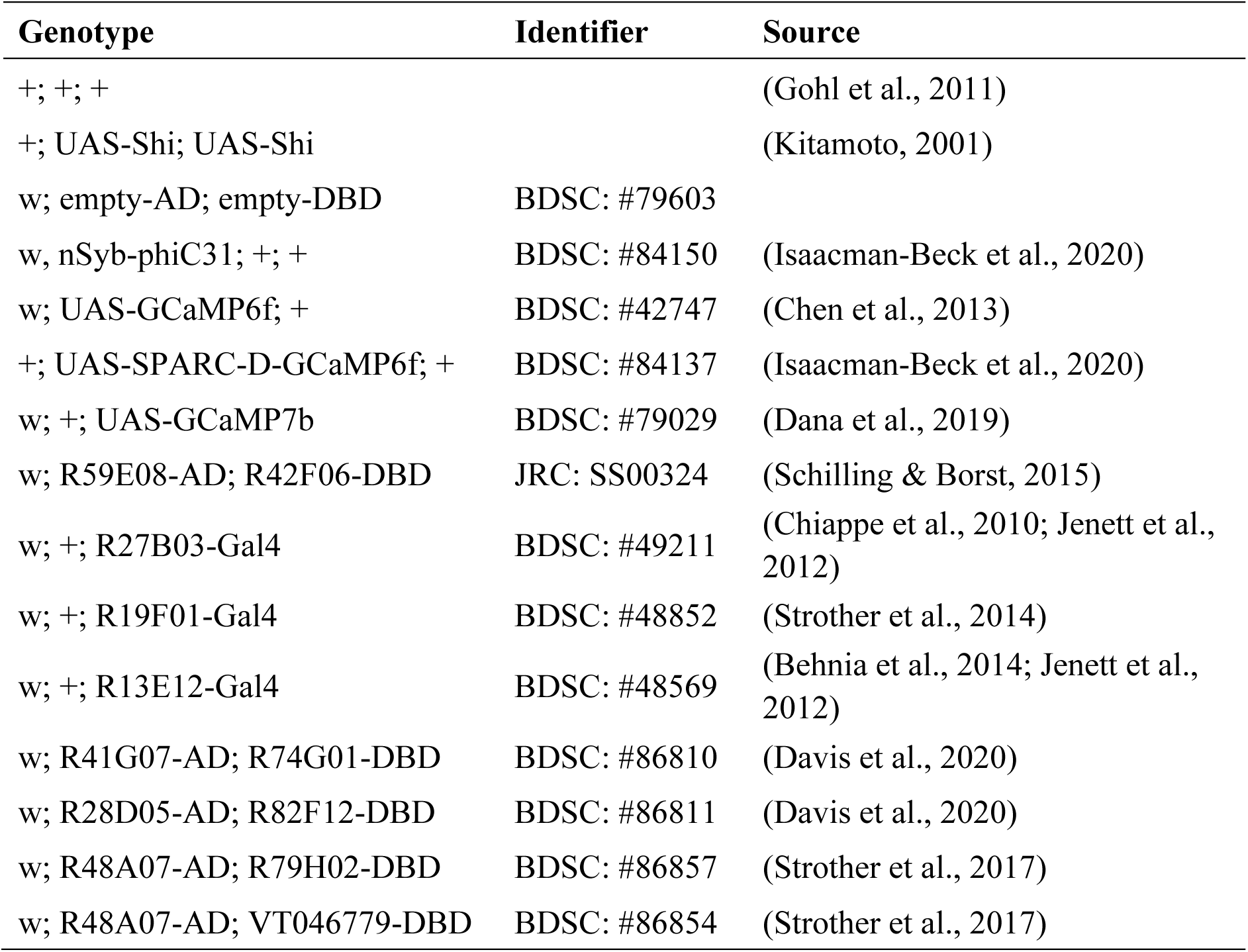

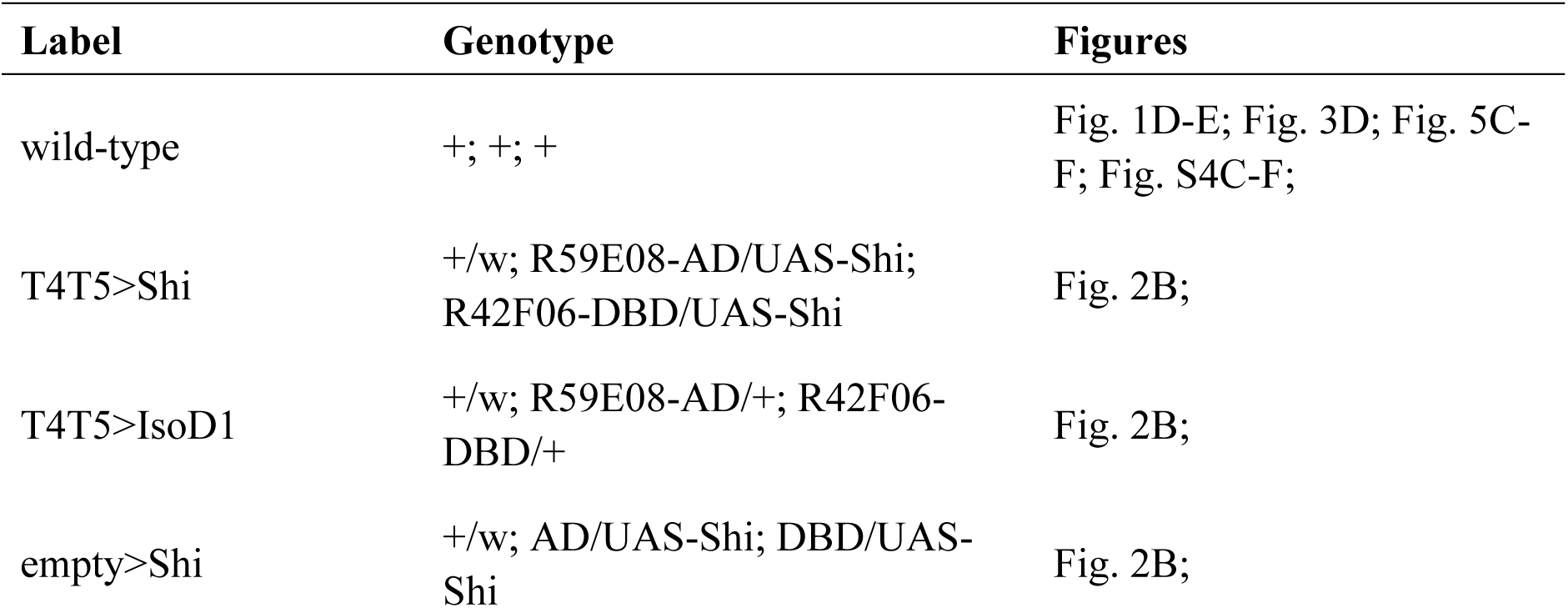
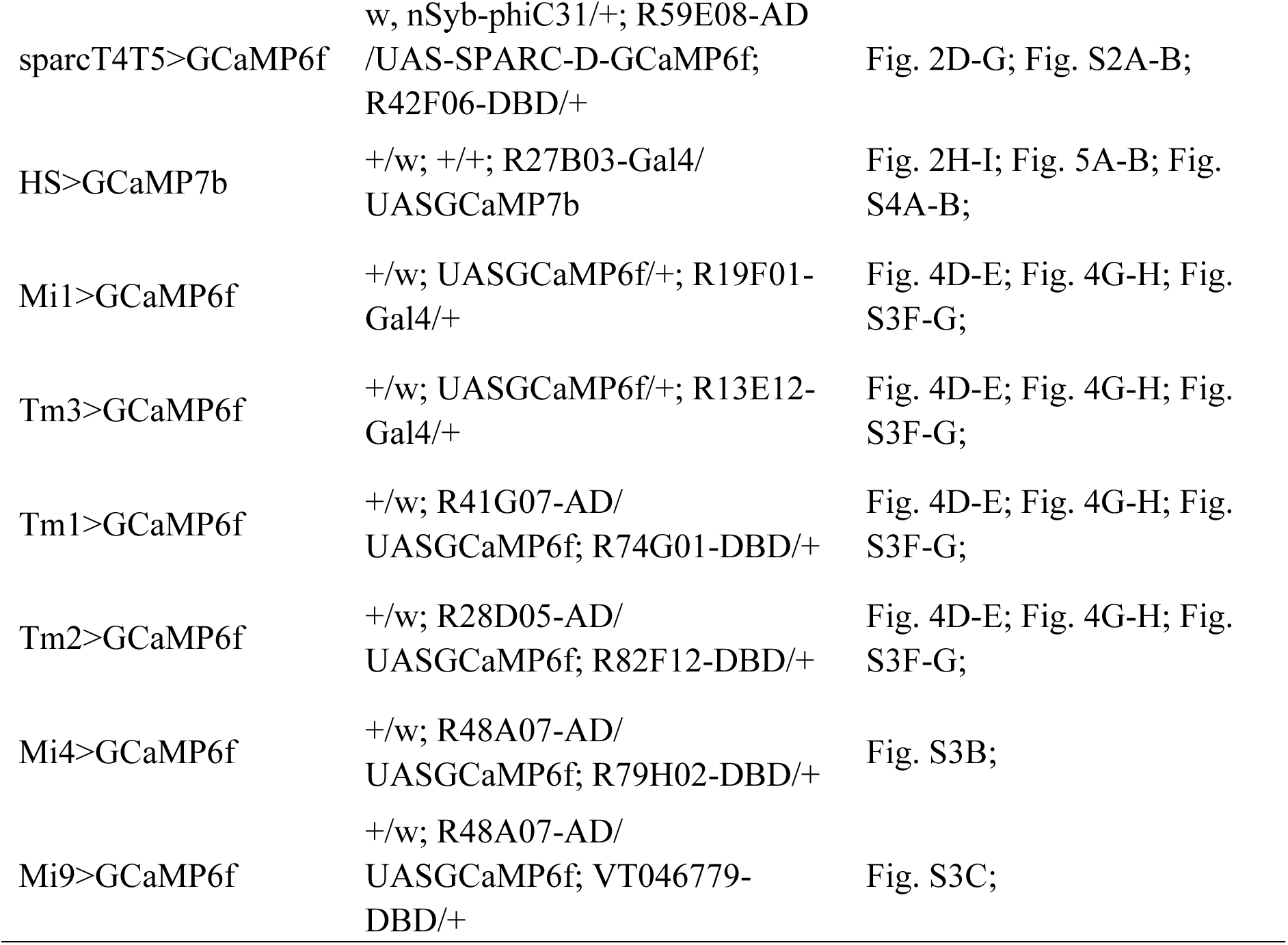

### Behavioral experimental setup and data analysis

Behavioral experiments followed procedures established in previous studies (Creamer et al., 2018). Flies were cold-anesthetized and tethered to surgical needles, then placed on top of air-supported balls. Visual stimuli were projected onto a surrounding panoramic screen, and fly walking responses were measured through ball rotations. The turning responses of flies during the stimulus presentation were averaged over time to compare average turning response. Temperature of the rig was set at 34-36°C to promote walking and activate thermogenetic tools.

### Experimental stimuli setup

Visual stimuli were generated using Matlab (Mathworks, Natick, MA, USA) and Psychtoolbox (Brainard, 1997; Kleiner et al., 2007; Pelli, 1997). Stimuli were presented on a virtual cylinder around the fly using a digital light projector (Texas Instruments, TX, USA) that refreshed the screen at 180 frames per second.

### Behavioral stimulus design

Sinusoidal interleave-shift stimuli: Full-contrast sinusoidal gratings with a spatial wavelength of 30° were presented in an alternating sequence with uniform full-field interleaves. Gratings were shown as a stationary image for 12/60 s, followed by an interleave with variable duration and contrast. Upon reappearance after the interleave, the gratings were spatially phase-shifted by a variable amount relative to its previous presentation, creating a stepwise motion pattern composed of discrete static gratings interrupted by uniform interleaves. Each stimulus epoch lasted 2s.

Checkerboard interleave-shift stimuli: Full-contrast random binary checkerboard gratings followed the same interleave-based presentation logic. Checks were 5° in azimuthal angle and full height vertically. A checkerboard grating was presented stationary for 16/60 s, then interrupted by a uniform full-field interleave with variable duration and contrast. Upon reappearance after the interleave, the checkerboard was spatially shifted by a variable amount. Each epoch lasted 2 s.

### Imaging experimental setup

Two-photon imaging followed protocols described in previous studies (Salazar-Gatzimas et al., 2016). Flies were cold-anesthetized and head-fixed to a custom metal holder using UV-cured epoxy. The cuticle and fat tissue covering the right optic lobe were surgically removed to expose right hemisphere of the brain, which was then bathed in oxygenated sugar-saline solution. The fly and holder were positioned above a panoramic screen with geometry similar to the behavioral experiments (Creamer et al., 2019). Imaging was performed using a two-photon microscope (Scientifica, Clarksburg, NJ, USA). Fluorescence signals were collected through a photomultiplier tube (PMT) and filtered with a 512/25 and a 514/30 bandpass filter (center/FWHM; Semrock, Rochester, NY, USA) to block stimulus light. Excitation was provided by a femtosecond Ti:sapphire laser (Mai Tai eHP, Spectra-Physics, Milpitas, CA, USA) tuned to 930 nm. Images were acquired at ∼13 Hz using ScanImage 5 software (Pologruto et al., 2003) and were subsequently motion-corrected offline. Before testing the neuronal response to the experimental stimulus, a probe specific to each neuron type was displayed for identifying responsive ROIs; the specific probe used is listed below in data analysis for each neuron type.

### Imaging stimulus design

Sinusoidal interleave-shift stimuli: Sinusoidal interleave-shift stimuli for imaging experiments are the same with behavioral ones except each stimulus epoch lasted 5s.

Sinusoidal pattern-uniform stimuli: Full-contrast sinusoidal gratings with a spatial wavelength of 30° were presented stationary for 12/60 seconds and then replaced by a uniform full-field interleave with variable contrast. The interleave lasted 2 seconds and was then terminated with a return to mean luminance.

### Imaging Data Analysis

T4/T5 neurons: Direction-selective T4 and T5 regions of interest (ROIs) were selected followed the criteria described in (Salazar-Gatzimas et al., 2016). Specifically, T4a, T4b, T5a, and T5b ROIs were identified based on their edge- and direction-selectivity. For each fly, ROIs were grouped such that for each stimulus condition, the GCaMP6f response to PD-shifted stimulus was defined as the T4a/T5a GCaMP6f responses to front-to-back-stimulus and T4b/T5b GCaMP6f responses to back-to-front-stimulus; responses to ND-shifted stimuli were constructed in the opposite manner. The averaged response of these ROIs was computed for each fly. Final results represent the mean responses across flies.

HS neurons: To identify HS neurons, we presented flies with visual probes consisting of square-wave gratings moving in the neuron’s preferred and null directions. ROIs were selected by computing the correlation between their GCaMP7b responses across four trials; only ROIs with an average inter-trial correlation greater than 0.4 were included. For each fly, responses from ROIs passing this threshold were averaged. Final results represent the mean responses across flies.

Medulla neurons: The selection of medulla neurons upstream of the direction-selective circuit followed methods described in prior work (Gonzalez-Suarez et al., 2022). Active ROIs were selected based on their responses to alternating light and dark flashes. To map azimuthal receptive field positions, we presented moving bars at a constant speed of 30°/s: bright bars for Mi1, Tm3, and Mi4; dark bars for Tm1, Tm2, and Mi9. The azimuthal receptive field position of each ROI was estimated from the timing of peak responses to these stimuli. ROIs from different flies of the same cell type were pooled. Each ROI was assigned a phase based on its receptive field position divided by the spatial wavelength of the grating. GCaMP6f responses were aligned by phase and replicated over two spatial cycles for better visualization.

### Motion energy simulation

All simulations were implemented in MATLAB. The simulation of a motion detector responding to interleave-shift stimulus followed a previous study (Zavatone-Veth et al., 2020), though note that this sort of linear-nonlinear model is inconsistent with some data (Badwan et al., 2019). We simulated a T4a motion detecting unit (ON pathway) in the right eye, which detects rightward motion. The T4a neuron receives excitatory input from upstream central neurons, and inhibitory input from flanking neurons on both sides (Arenz et al., 2017; Shinomiya et al., 2019; Takemura et al., 2017). The central neuron and the flanking neuron to its right are ON neurons, while the flanking neuron on the left side of the central neuron is an OFF neuron.

All 3 input signals were modeled with the identical Gaussian spatial filter:

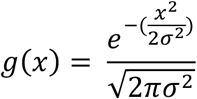

In the simulation, we used 𝜎 = 2.42°, corresponding to a full-width-half-max of 5.7° (Stavenga, 2003). This spatial filter neglects surround antagonism in these inputs to T4 (Arenz et al., 2017).

All signals were modeled with a high pass temporal filter:

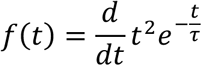

In the simulation, we used 𝜏 = 0.05 s, for the central signal, and 𝜏 = 0.1 s for the two flanking signals. The temporal filters were normalized to have unity L2 norm before being combined with the spatial filters.

Following prior modeling work (Zavatone-Veth et al., 2020), the spatiotemporal filters of the influence of inputs onto T4 are:

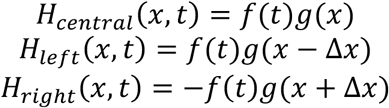

The spatiotemporal filter of the motion detector unit is:

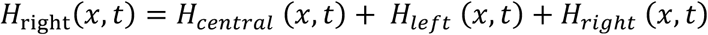

In the simulation, we used Δ𝑥 = 5°.

The stimulus 𝑆(𝑥, 𝑡) is the same sinusoid interleave-shift stimulus we used in the experiments. Sinusoidal gratings with a spatial wavelength of 30° were presented in an alternating sequence with uniform full-field interleaves. Each time, gratings were shown as a stationary image for 12/60 s, followed by an interleave with variable duration and contrast. Upon each reappearance, the gratings were spatially phase-shifted by ¼ cycle to the right relative to their previous presentation.

In the simulation, we used interleave durations of 𝑇_0+’_ = {0, 1, 2, 4, 8, 12, 16, 20}/60𝑠 with zero contrast interleave, and interleave contrast 𝑐_0+’_ = {−1, 0, 1} with 12/60s duration.

The filter weights of this rightward motion detector across spatial frequency (k) and temporal frequency (ω) were obtained by applying a two-dimensional Fourier transform to the spatiotemporal filter of the rightward motion detector 𝐻(𝑥, 𝑡):

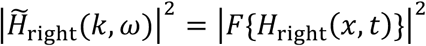

Its counterpart was spatially flipped to yield right-left opponency:

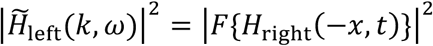

The filter weights of a full opponent motion detector were obtained by subtraction of the rightward selective filter and its leftward selective opponency:

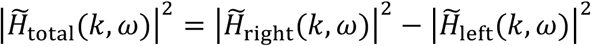

The motion energy of stimulus across 𝑘 and 𝜔 were obtained by applying a two-dimensional Fourier transform to 𝑆(𝑥, 𝑡):

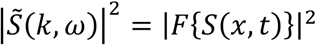

For each stimulus, the spatiotemporal mean motion energy (𝑅^K^) of different directions was obtained by multiplying the filter weight and energy at their corresponding 𝑘 and 𝜔 and averaging over frequencies (Adelson & Bergen, 1985):

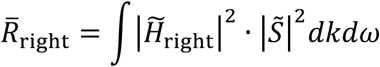

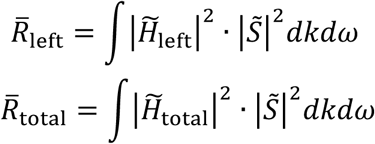

